# Gradual differentiation uncoupled from cell cycle exit generates heterogeneity in the epidermal stem cell layer

**DOI:** 10.1101/2021.01.07.425777

**Authors:** Katie Cockburn, Karl Annusver, Smirthy Ganesan, Kailin R. Mesa, Kyogo Kawaguchi, Maria Kasper, Valentina Greco

## Abstract

High turnover tissues continually lose specialized cells that are replaced by stem cell activity. In the adult mammalian epidermis, it is unclear how molecularly heterogenous stem/progenitor cell populations fit into the complete trajectory of epidermal differentiation. We show that differentiation, from commitment to exit from the stem cell layer, is a multi-day process wherein cells transit through a continuum of transcriptional changes. Differentiation-committed cells remain capable of dividing to produce daughter cells fated to further differentiate, demonstrating that differentiation is uncoupled from cell cycle exit. These cell divisions are not required as part of an obligate transit amplifying program but instead protect density in the stem cell layer. Thus, instead of distinct contributions from multiple progenitors, a continuous gradual differentiation process fuels homeostatic epidermal turnover.

**One sentence summary:** Heterogeneity in the epidermal stem cell layer reflects a gradual differentiation program that is uncoupled from the loss of proliferative capacity.

---

Highly regenerative tissues such as the skin, blood and intestine are under continuous cell turnover, relentlessly producing differentiated cells to replace those that are lost. How such tissues orchestrate the complex transition from undifferentiated stem cell populations towards post-mitotic, molecularly distinct and often spatially segregated differentiated cell types is not fully understood. Although cells at intermediate stages of this transition, often termed transit amplifying cells, are hypothesized to play a necessary role in fueling tissue turnover (*1*–*3*), how such cell states arise and whether they are essential for homeostasis in many cases remains unclear.

In the stratified mammalian skin epidermis, cells from an underlying, highly proliferative basal layer differentiate and move upwards to replace outer, barrier-forming cells that are shed on a daily basis **(Figure S1A)**. Extensive efforts have been made to define the cell states and correspondent behaviors within the basal layer that fuel this lifelong tissue turnover. Several studies support a model of epidermal homeostasis through the cooperative efforts of multiple, molecularly distinct progenitor types **(Figure S1B)** that include fast-cycling, differentiation-primed committed progenitors underpinned by a second, molecularly distinct slow-cycling stem cell population (*4*, *5*), pairs of directly coupled stem and differentiation-committed progenitor cells (*6*), or two independent stem cell populations with different cycling kinetics and transcriptional profiles (*7*). In contrast, other studies instead propose a single type of proliferating progenitor population that generates dividing and differentiating cells with equal probability at the population level (*8*–*11*) **(Figure S1B)**. It is not clear how the progenitor populations that have been proposed fit into the epidermal differentiation journey itself. Specifically, we do not know when differentiation begins, how long it lasts, and how the real-time behaviors of individual cells correspond to expression of known molecular markers.

To understand this journey in more detail, we focused on the long-standing observation that a subset of basal cells express the well-established differentiation marker Keratin 10 (K10) (*9*, *12*, *13*). Because these cells have typically been considered a post-mitotic population in the process of exiting the basal layer, neither their real-time behaviors nor their relevance to any of the aforementioned models has been closely examined. To characterize K10-expressing basal cells in greater detail, we performed whole-mount immunostaining **(Figure 1A)**. Within all basal cells, 43% stained positive for K10, with the majority of these K10 positive cells making a small area of contact (footprint) with the underlying extracellular matrix (ECM) as would be expected from delaminating basal cells **(Figure 1A, B)**. Surprisingly however, we also observed K10 positive cells with a typical basal-cell morphology including a normal-sized ECM-footprint **(Figure 1A, B)**. These results indicate that in contrast to previous models (*14*), the differentiation process of basal epidermal cells may begin even prior to a detectable change in cell morphology.

**Figure 1.**
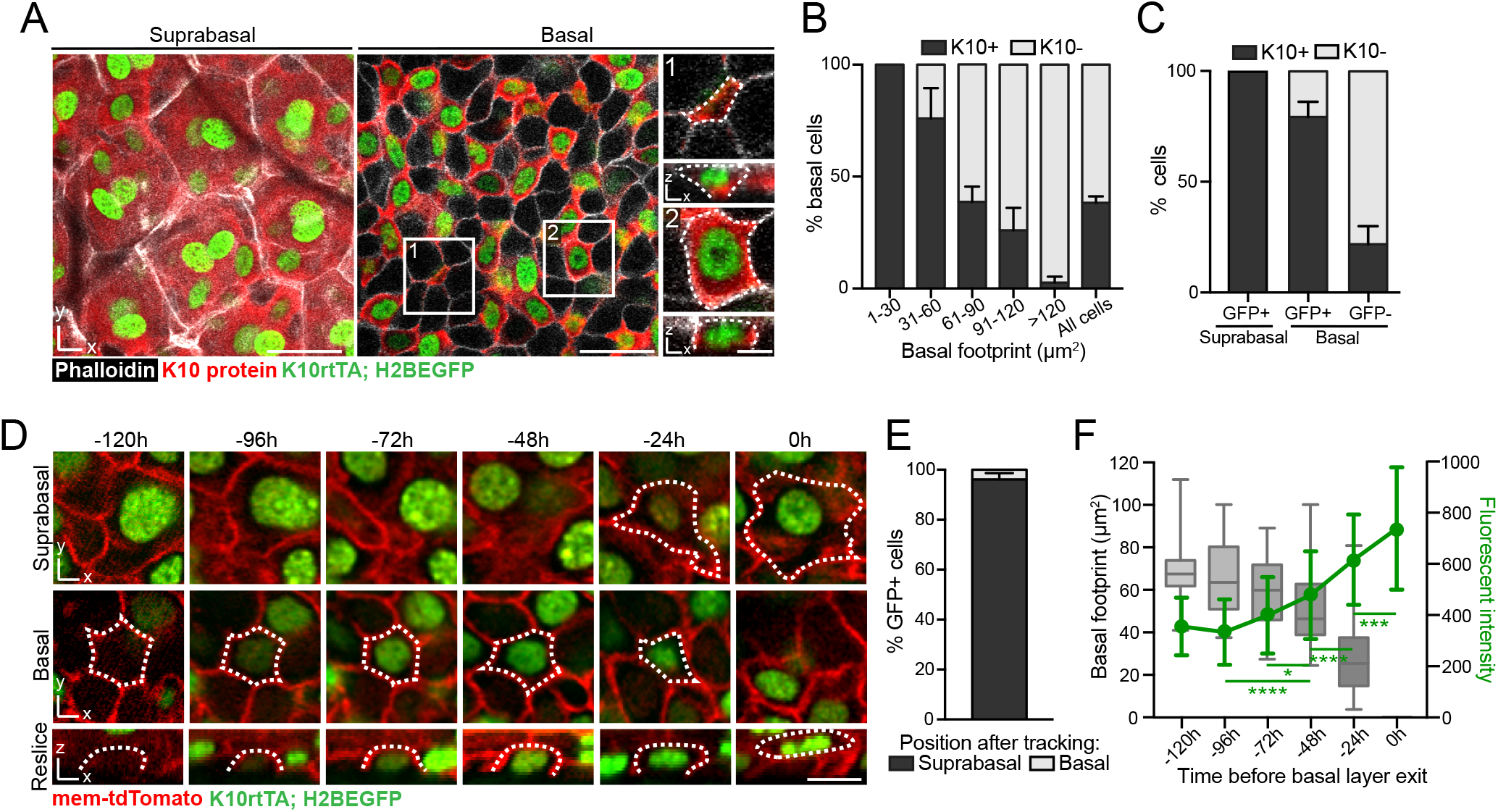
Epidermal stem cell differentiation occurs over multiple days. **(A)** Representative whole mount staining showing Keratin 10 protein (red) and K10 reporter (*K10rtTA; pTRE-H2BGFP*) expression (green) in suprabasal and basal cells. Cell boundaries are visualized with phalloidin (white). Insets 1 and 2 show examples of Keratin 10-positive basal cells with very little (1) or average (2) amounts of ECM contact. Scale bar=25μm (large field of view) or 5μm (insets). **(B)** Quantification of percent basal cells with the indicated levels of ECM contact that also express Keratin 10 protein. N=500 cells from 3 mice. **(C)** Quantification of the overlap between Keratin 10 protein expression and K10 reporter expression in basal and suprabasal cells. N=800 cells from 3 mice. **(D)** Representative images of a revisited basal cell as it induces K10 reporter expression and later exits the basal layer. Scale bar=10μm. **(E)** Quantification of basal vs. suprabasal position in basal cells scored as K10 reporter positive on Day 0 and revisited for up to 10 subsequent days. N=127 cells from 3 mice. **(F)** Quantification of K10 reporter expression (green line) and basal cell-ECM contact (box and whisker plot) in the days preceding basal layer exit. N=74 cells from 2 mice. ANOVA, p<0.0001; Tukey’s HSD, p<0.0001 (96h vs 48h and 48h vs 24h), p<0.001 (24h vs 0h), and p<0.05 (72h vs 48h). For box and whisker plots in (F), box boundaries represent 25^th^ and 75^th^ percentiles, and error bars represent max and min values. For bar graphs in (B), (C) and (E) and line graph in (F), error bars represent S.D.

To understand the temporal dynamics of K10 expression in living basal cells, we performed intravital imaging (*10*, *15*) using a reporter system in which the *Krt10* promoter drives H2BGFP fluorescence (*K10rtTA*; *pTRE-H2BGFP*) (*16*) **(Figure 1A, C)**. By combining this reporter with a ubiquitous plasma membrane marker (unrecombined *mTmG*) and revisiting the same epidermal regions every 24h **(Figure 1D)**, we found that the vast majority (96%) of K10 reporter positive basal cells could be seen exiting the basal layer in subsequent timepoints **(Figure 1E)**, indicating that this population has largely committed to differentiate. To visualize this journey in its entirety, we then focused on reporter negative cells that could be seen inducing H2BGFP expression during our revisits. Notably, K10 reporter signal almost always became visible 1-2 days before cells began to exhibit the first morphological signs of exiting the basal layer, and this delamination process occurred surprisingly slowly over a further average of 36h **(Figure 1D, F and S1C-E)**. In contrast to the rapid, actomyosin-based extrusion events that have been described in embryonic epidermis and other systems (*17*–*19*), basal cell delamination lacked obvious signs of ring-like actin or myosin accumulation **(Figure S1F, G)**. Thus on average, newly positive cells could be seen progressively increasing their H2BGFP signal for approximately 4 days before basal layer exit **(Figure 1D, F)**. These temporal observations demonstrate that adult basal cell differentiation, from commitment to the completion of delamination, is a gradual multi-day process.

We next aimed to understand how K10 expression relates to the global transcriptional changes associated with basal cell differentiation and to other differentiation-primed progenitor populations that have been described (*4*–*6*). Our previous reconstruction of the epidermal differentiation trajectory, based on single-cell transcriptomes of randomly sampled cells from basal and suprabasal layers, grouped the cells according to their individual gene expression from basal (*Krt14*^high^), mature (*Krt*10^high^) to terminally differentiated (*Lor*^high^) cell states (*20*). To define the basal-suprabasal border on this trajectory, we generated single-cell transcriptomes of FACS-isolated basal cells (Live/ITGA6+/SCA1+/CD34−; **Figure S2A**), which we merged with two published datasets that include both basal and suprabasal epidermal cells **(Figure S2B)**. After computational removal of cells from other epithelial compartments (Methods), the combined dataset allowed us to assign the basal-suprabasal border according to the sorted basal (ITGA6+) cells **(Figure 2A, Methods)**. Notably, the previously defined intermediate *Krt10*^dim^/*Ptgs1*^dim^/*Mt4*^+^ cell group (Differentiated I), as well as some cells of the mature *Krt10*^high^/*Ptgs1*^high^ group (Differentiated II) (*20*), are basal-layer cells **(Figure S2C-D)**. This is exemplified by ~40% of basal cells already expressing *Krt10* **(Figure S3A-B)**, as well as expression of basal and differentiation genes in the same cells **(Figure 2B-C, S3C)**, consistent with recent studies in skin as well as oral epithelia (Figure S3D)(*6*, *21*, *22*). To more closely examine where the first transcriptional changes begin (signifying the onset of differentiation), we grouped the cells along the trajectory into 10 differentiation bins **(Figure 2D, S3E, Methods)**, revealing *Krt10* as the first upregulated differentiation marker (bin 3), followed by *Krtdap* and *Krt1* (bin 4), *Mt4* and *Sbsn* (bin 5), and *Ivl* and *Lor* (bin 6) **(Figure 2D, S3F-H)**. Basal marker-gene expression also starts to decrease in bin 3 (*Itga6, Ly6a*), followed by a marked decrease of *Krt14* in bin 4 **(Figure 2D, S3F-H)**. Thus the earliest molecular changes associated with differentiation start in bin 3, marked by an increase of *Krt10* expression in cells displaying otherwise typical characteristics of stem cells (*Krt14*^high^, large ECM-footprint) **(Figure S3C, arrowhead)**. These results indicate that instead of discrete intermediate cell states, basal cells differentiate through a series of progressive transcriptional changes, raising questions about how previously proposed differentiation-primed progenitor populations, most notably those that can be traced with involucrin CreER (*4*, *5*), fit within this continuum.

**Figure 2.**
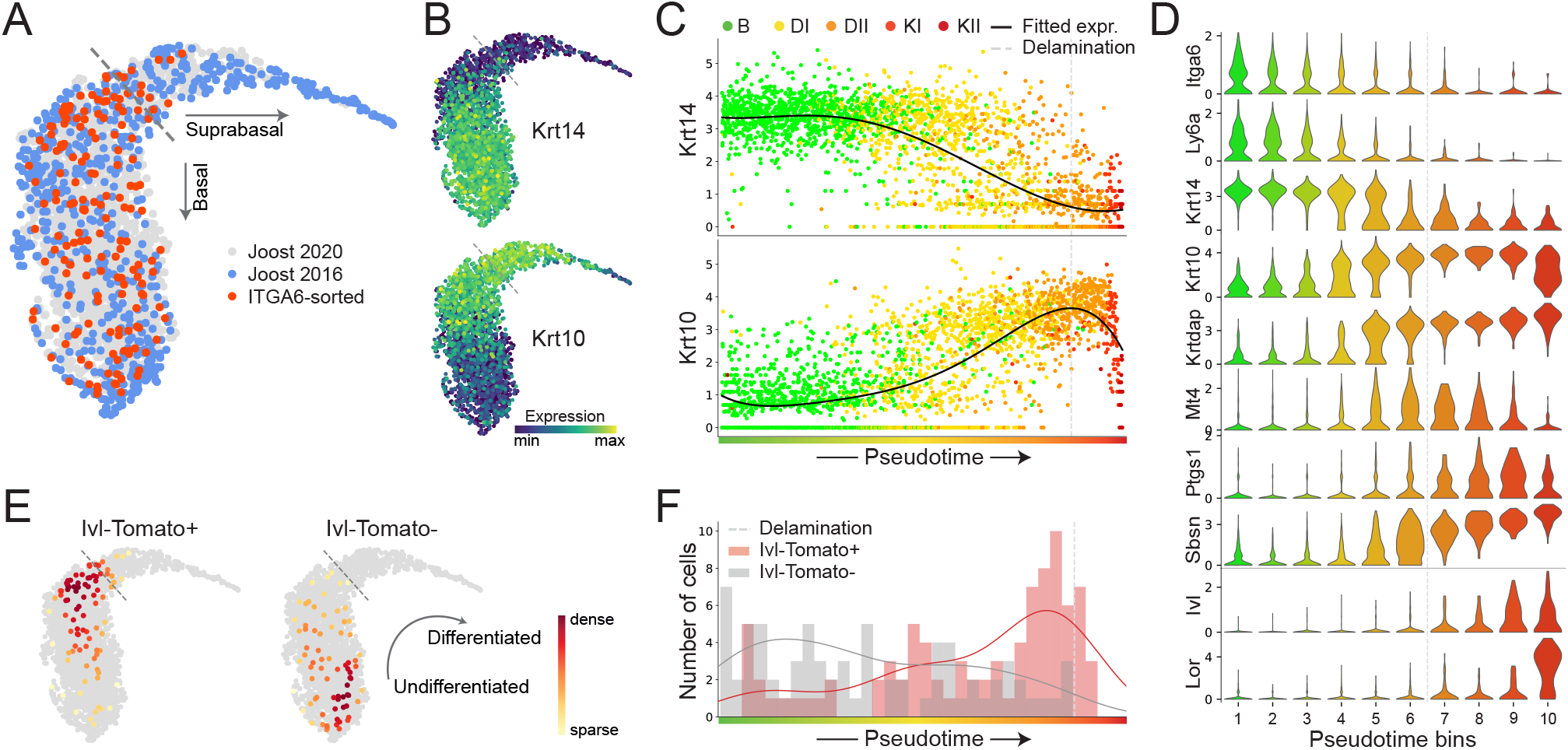
Single-cell RNA-seq trajectory analysis of epidermal stem cell and committed progenitor differentiation. **(A)** UMAP of the combined epidermal IFE datasets, showing the distributions for Joost 2020, Joost 2016 and ITGA6-sorted cells. **(B)** *Krt14* and *Krt10* gene expression patterns overlaid on the combined UMAP. **(C)** *Krt14* and *Krt10* expression changes for individual cells, as well as fitted expression, ordered along pseudotime and colored according to their differentiation state from basal (B), differentiated (DI and DII) to keratinized (KI and KII). **(D)** Violinplots of differentiation-associated gene expression within cells grouped into 10 pseudotime bins. **(E)** Location of *Ivl*-traced (Tom+) and non-traced (Tom-negative) sorted cells on the combined UMAP. Cells are colored according to the local density of visualized populations. **(F)** Distribution of *Ivl*-traced and non-traced cells along the pseudotime (histogram), together with estimation lines for cell density. (A-F) Dashed lines indicate the assigned delamination point based on the location of 95% of ITGA6-sorted cells (Methods). (B-D) Expression is shown as log-normalized counts.

We focused on involucrin CreER traced cells by acquiring our ITGA6-sorted dataset from 2-day-traced *IvlCreERT2; R26-Tomato* mice, with separately sorted and sequenced committed progenitor (Tom+) and stem cells (Tom-negative) **(Figure S2A)**. While these traced and non-traced cells could be found spread out across the pseudotime, the majority of Tom+ basal cells indeed mapped with the *Krt10*+/*Mt4*+ cells and the majority of the Tom-negative cells mapped with basal *Krt14*+ cells **(Figure 2E, F, S3I-J)**, suggesting that the traced cell population largely represents cells that are committed to differentiation but not necessarily a discrete progenitor cell state. Further comparison of individual cells along the differentiation trajectory revealed that *Krt10*-expressing cells arise before *Ivl*-traced cells during basal cell differentiation **(Figure 2C, F)**. Therefore, the analysis of K10 positive basal cell behaviors is likely to reveal cells transitioning through other states of differentiation that have previously been described. In sum, this fine-tuned differentiation trajectory appoints *Krt10* expression at the molecular onset of a continuum of transcriptional changes associated with epidermal differentiation.

To our surprise, further analysis of the cell cycle stages within our dataset revealed that approximately 24% of S/G2/M phase cells express *Krt10* mRNA **(Figure3A, B and S4A-C)** and staining for K10 protein was detectable in approximately 14-18% of S/G2/M phase cells **(Figures 3C, D and S4D, E)**. To examine these events in more detail we performed timelapse imaging and were able to observe K10 reporter positive cells undergoing mitosis **(Figure 3E)**. We found that these divisions occurred parallel to the basement membrane, producing daughter cells that were fully integrated within the basal layer and retained reporter expression **(Figure 3E and Movie S1)**. To uncover the consequences of these K10 positive proliferation events, we tracked the resulting daughter cells and quantified their fates. In the 3 days following division, most daughter cells (60%) had delaminated, completing the differentiation trajectory begun by their mother cell **(Figure 3F, G)**. More rarely (6% of cases), daughter cells underwent an additional round of division. In cases when daughters from these subsequent divisions could be further tracked, we witnessed them also delaminating **(Figure S4F)**. These results contrast with K10 negative divisions, where 43% of daughters had divided and 20% had delaminated after 3 days **(Figure 3G)**.

**Figure 3.**
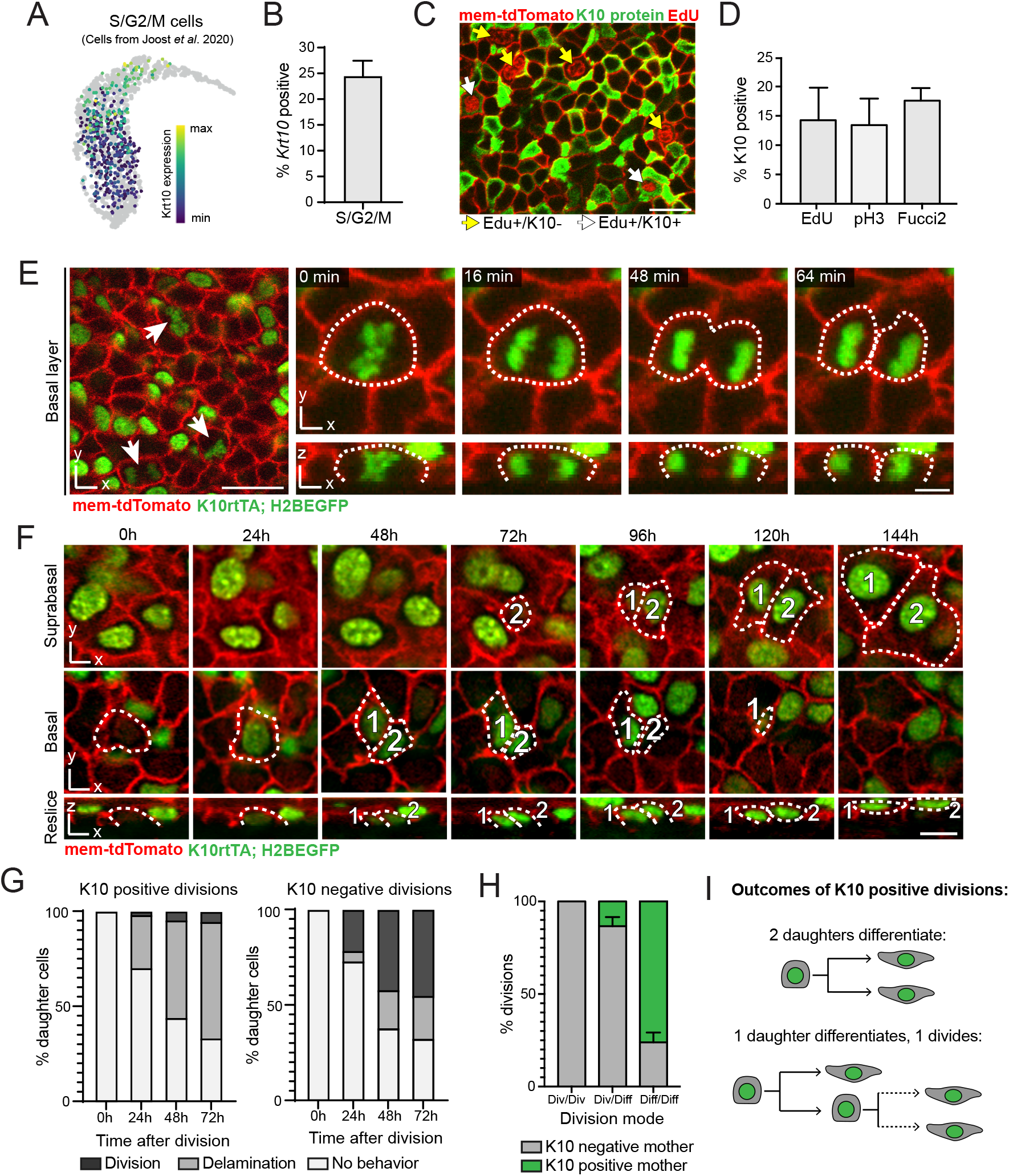
Differentiation-committed cells proliferate during homeostasis. **(A)** UMAP showing *Krt10* gene expression levels in all cycling (S and G2/M) epidermal cells from the Joost 2020 dataset. **(B)** Quantification of *Krt10*-positive cells (cutoff=1.84, see Methods) within the proliferative cell population (S/G2/M) from the Joost 2020 dataset. N=5 mice. **(C)** Representative whole mount staining of EdU incorporation (red nuclei), K10 (green), and mem-tdTomato (red membrane) showing both EdU-positive, K10-negative (yellow arrows) and EdU-positive, K10-positive (white arrows) basal cells. Scale bar=20μm. **(D)** Quantification of percent cycling cells, represented by EdU, pH3, or Fucci2 mVenus-hGem positivity, that are classified as K10 protein positive. N=199 cells from 2 mice, 507 cells from 3 mice, and 484 from 3 mice for EdU, pH3, and Fucci2 signal respectively. Error bars indicate S.D **(E)** Single timepoints (left) or stills from timelapse imaging (right) show K10 reporter positive (green) mitotic figures, indicated by dotted lines. Timelapse imaging captures K10 reporter positive divisions generating two basal daughter cells. Membrane is visualized with mem-tdTomato (red). Scale bar=20μm (large field of view) or 5μm (timelapse stills). **(F)** Representative images of a revisited basal cell as it induces K10 reporter expression and divides to produce two daughter cells (numbered 1 and 2) that exit the basal layer. Scale bar=10μm. **(G)** Quantification of cumulative daughter fates in the 3 days following K10 reporter positive and K10 reporter negative divisions. N= 114 cells from 3 mice (K10 reporter positive divisions) and 186 cells from 3 mice (K10 reporter negative divisions). **(H)** Quantification of division modes (asymmetric; symmetric with both daughters differentiating; or symmetric with both daugthers dividing) from all division events where the subsequent behavior of both daughter could be resolved in later revisits. N= 127 divisions from 3 mice **(I)** Schematic of daughter cell fates after K10 reporter positive divisions. For bar graphs in (B), (D) and (H), error bars represent S.D.

As homeostatic basal cell divisions can lead to asymmetric or symmetric fate outcomes (one daughter divides and one differentiates, or both daugthers perform the same behavior, respectively) (*4*, *8*), we wanted to understand how K10 reporter positive divisions contribute to these different modes of proliferation. Thus we focused on all cell division events where the behavior of both daughter cells could be resolved in subsequent days of imaging. We found that 76% of all symmetric divisions that produce two differentiating daughters came from K10 positive mother cells, while only 18% of asymmetric divisions and no symmetric divisions that produce two dividing daugthers had K10 positive mothers **(Figure 3H, I)**. This preponderance of symmetric, differentiation-fated divisions contrasts with the largely asymmetric, self-renewing mode of division that has been proposed to characterize differentiation-primed progenitors in other models (*4*, *5*). Together these results indicate that, although K10 reporter expression signifies commitment to eventually delaminate, differentiating cells remain capable of dividing to generate short-term lineages of basal daughter cells fated to exit the stem cell layer.

We next sought to understand the factors driving the division of differentiating cells. One possibility is that just like their undifferentiated neighbors, differentiating cells divide as a consequence of basal layer density changes when nearby cells are lost to delamination (*23*). Conversly, these divisions may occur irrespective of neighbor delaminations, driven instead by an intrinsic program of amplification **(Figure 4A)**. To distinguish between these two possibilities, we quantified the behaviors taking place within 10μm (a one-cell distance) in the days leading up to K10 reporter positive division events. If these divisions occur in response to a neighboring delamination, they should be preceded by the net loss of one nearby cell. If K10 divisions occur in response to other cues, this imbalance will not be observed **(Figure 4A)**. Notably, a clear net loss of one neighbor preceded K10 reporter positive divisions, just like those of K10 negative cells, indicating that differentiating basal cells indeed divide as a response to loss of delaminating cells in their local neighborhood **(Figures 4B)**.

**Figure 4.**
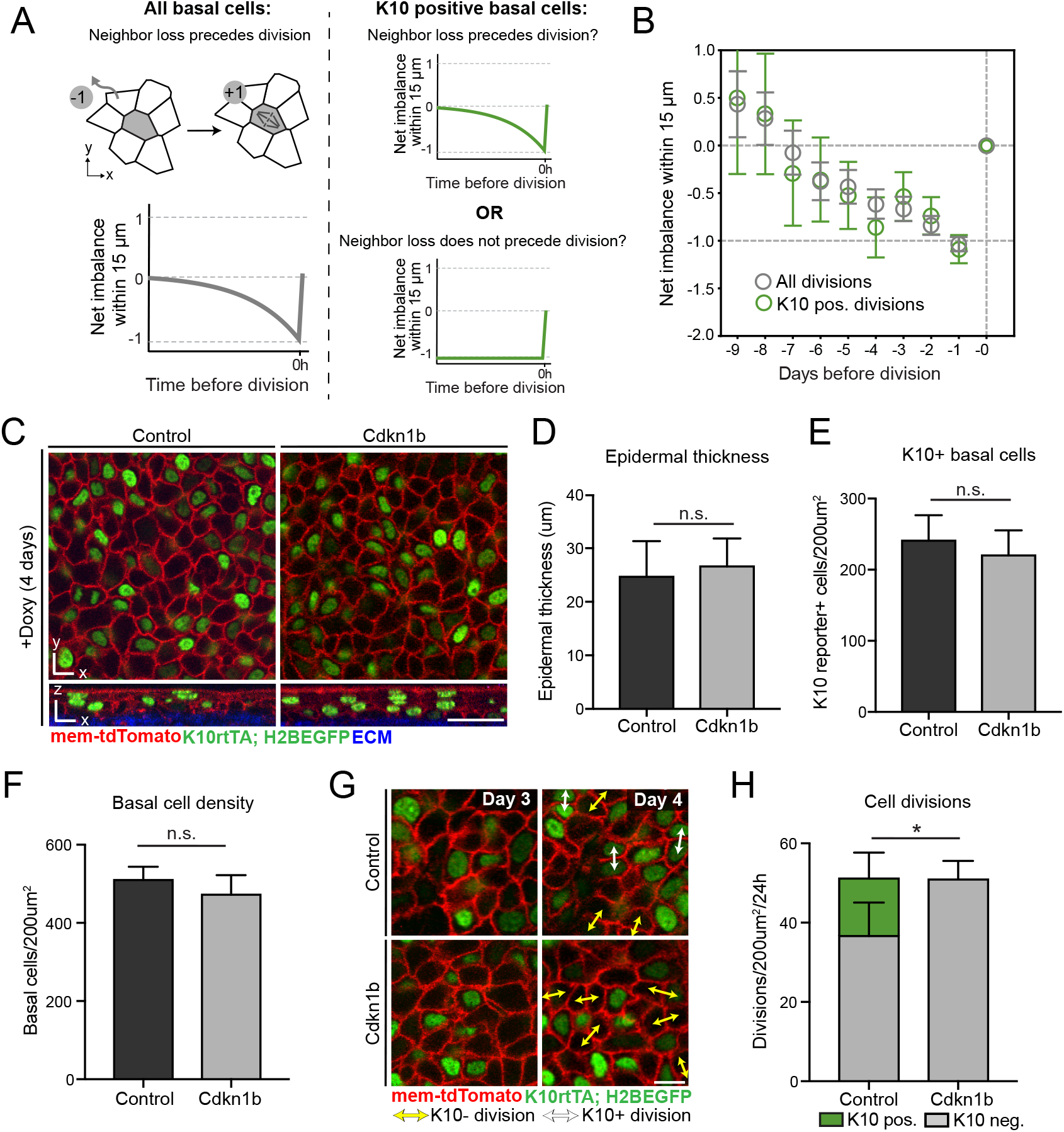
Division of differentiating cells is not required for epidermal maintenance. **(A)** Schematic of possible neighbor imbalance scenarios in the days leading up to K10 reporter positive divisions. Cumulative neighbor loss through delamination (−1) and neighbor gain through division (+1) was scored in the days leading up to division events (see Methods for details). If K10 reporter positive cells, like the basal population as a whole (left panel), proliferate in response to neighbor loss, imbalance will drop to −1 before division (right panel, top green line). If K10 reporter positive cells are unaffected by neighbor loss, no imbalance will occur before division (right panel, bottom line). **(B)** Quantification of fate imbalance leading up to division events. N=262 reporter negative and 68 reporter positive cells from 2 mice. **(C)** Representative images of Control (*K10rtTA; pTRE-H2BGFP*) and Cdkn1b (*K10rtTA; pTRE-H2BGFP; pTRE-Cdkn1b*) epidermis after 4 days of Doxycycline administration. Top row: xy section of basal layer. Bottom row: lateral reslice. K10 reporter is shown in green and membrane is visualized with mem-tdTomato (red). Scale bar=25μm. **(D)** Quantification of total epidermal thickness in Control and Cdkn1b mice after 4 days of Doxycycline administration. Student’s t-test, p>0.05. **(E)** Quantification of total K10 reporter positive cells in the basal layer of Control and Cdkn1b mice after 4 days of Doxycycline administration. Student’s t-test, p>0.05. **(F)** Quantification of basal density in Control and Cdkn1b mice after 4 days of Doxycycline administration. Student’s t-test, p>0.05. **(G)** Representative images of Control and Cdkn1b basal cells between Day 3 and Day 4 of Doxycycline administration, showing K10 reporter negative divisions (yellow arrows) and K10 reporter positive division (white arrows). Membrane is visualized with mem-tdTomato (red). Scale bar=10μm. **(H)** Quantification of K10 reporter positive and K10 reporter negative divisions in Control and Cdkn1b mice between Day 3 and Day 4 of Doxycycline administration. Student’s t-test comparing K10 reporter negative divisions (gray bars), p<0.05. For (D), (E), (F) and (H), N= 2×200μm^2^ regions per mouse from at least 3 mice per genotype, and error bars represent S.D.

The proliferation of differentiation-primed progenitors has been hypothesized to play an essential role in epidermal homeostasis (*4*, *24*), but the necessity of this behavior has never been directly tested. To evaluate the role of K10 positive divisions during epidermal homeostasis, we used the *Krt10* promoter to induce the cell cycle inhibitor *Cdkn1b* (or p27) (*25*) in differentiating cells and monitored these cells with the K10 reporter (*K10rtTA*; *pTRE-Cdkn1b; pTRE-H2BGFP*) **(Figure S5A)**. Within 24h of doxycyline treatment we observed a complete loss of K10 reporter positive mitotic cells within the basal layer, indicating a rapid and penetrant block of proliferation within this population **(Figure S5B)**. We then assessed potential consequences to the differentiation process in the days directly following Cdkn1b induction. To understand whether the number of suprabasal layers was altered, we measured total epidermal thickness and found it to be unchanged **(Figure 4C, D)**. In agreement with this observation, the number of cells delaminating out of the basal layer, as measured by complete loss of contact with the ECM, was comparable in tissue with and without Cdkn1b induction, demonstrating that cell division is not required for the later maturation or movement of differentiating cells out of the basal layer **(Figure S5C)**. Most interestingly, we found that expression of Cdkn1b did not affect the number of K10 reporter positive basal cells **(Figure 4E)**, indicating that the size of the differentiating cell pool is maintained even when these cells are entirely unable to divide. Thus proliferation of the K10 positive population is not required to generate the differentiated cells needed to sustain regeneration.

Given that K10 reporter positive divisions make up approximately one quarter of all proliferation events in the basal layer **(Figure 4H)**, we were surprised to observe that the basal cell density remained unaffected even after 4 days of Cdkn1b induction **(Figure 4F)**. To understand how this maintenance was achieved, we directly tracked the number of cell divisions taking place in Cdkn1b versus control tissue over a 24h period and found that K10 negative basal cells in the Cdkn1b epidermis increased their proliferation rate to equal the number of divisions performed by both K10 positive and K10 negative cells in control tissue **(Figure 4G, H)**. These results indicate that the need for proliferation in the homeostatic basal layer is normally satisfied by the contribution of both differentiating and undifferentiated cells. Thus, proliferation of differentiating cells in the adult epidermis occurs as a consequence of the need to replace delaminating neighbors while still in the basal layer, and not as part of an obligate transit amplifying program that fuels proper numbers of differentiating cells.

In summary, this study defines a gradual and progressive differentiation process that reconciles seemingly disparate models of the cell populations and behaviors that sustain homeostasis of the adult epidermis **(Figure S5D)**. We provide evidence that molecularly heterogeneous cell states in the basal layer (*4*, *6*, *13*) do not represent distinct populations of self-renewing progenitors, but instead can be seen as snapshots of cells at different points along a single continuum of differentiation that is carried out over several days. Since proliferative capacity is not lost until well after these changes are initiated, differentiating cells can produce short-lived lineages of daughter cells fated to exit the stem cell layer, an observation not predicted by single progenitor models (*8*–*10*). Strikingly, division of differentiation-committed cells helps to replace basal neighbors that are lost to differentiation, but these events are not necessary for the differentiation process itself or for overall maintenance of the tissue. We propose that instead of distinct contributions from multiple progenitor types, a single continuous differentiation process fuels the lifelong turnover of the epidermis.

## Supporting information

Cockburn et al_supplemental materials

Cockburn et al_Movie S1

## Acknowledgements

We thank all members of the Greco and Kasper labs for critical feedback on the manuscript. We thank T. Lechler for *K10-rtTA* mice, E. Fuchs for *pTRE-H2BGFP* mice, R. Weigert for *Lifeact-GFP* mice, R. Adelstein for *GFP-NMMIIB* mice and S. Aizawa for *R26p-Fucci2* mice.

## Author contributions

K.C., K.A., K.K., M.K. and V.G designed experiments and analyzed data. K.C. performed 2-photon imaging, whole mount staining, mouse genetics and image analysis. K.A. performed smRNA-FISH and single-cell RNA-sequencing experiments. K.A. and M.K. performed single-cell transcriptome analysis and interpretation. K.K. performed data analysis and statistical modeling. S.G. assisted with whole mounts and mouse genetics. K.M. performed time-lapse imaging. K.C. and V.G. wrote the manuscript with input throughout from K.A., M.K., S.G. and K.K.

## Data availability

All data from this study are available from the authors on request. The accession number for the sequencing data reported in this paper is GEO: GSE152044. The complete computational analysis workflow for the scRNAseq data will be available in the form of jupyter notebooks at https://github.com/kasperlab. The python scripts for the image analysis will be available on request.

