## Supplementary material for "Gradual differentiation uncoupled from cell cycle exit generates heterogeneity in the epidermal stem cell layer": Cockburn et al_supplemental materials

**This PDF file includes:**

- Materials and Methods

- Figs. S1 to S6

- Captions for Movie S1

**Other Supplementary Materials for this manuscript includes the following:**

- Movie S1

**Materials and Methods:**

**Mice and experimental treatment**

*mTmG* (*26*), *K14-CreER* (*27*), *tetO-Cdkn1b* (*25*), *Ivl-CreERT2* (*28*) and *R26-tdTomato* (*29*) mice were obtained from the Jackson Laboratory. *K10-rtTA*(*16*) mice were obtained from T. Lechler (Duke University), *pTRE-H2BGFP* (*30*) mice were obtained from E. Fuchs (Rockefeller University), *Lifeact-GFP* (*31*) mice were obtained from R. Weigert (NIDCR, NIH), *GFP-NMMIIB* (*32*) mice were obtained from R. Adelstein (NHLBI, NIH) and *R26p-Fucci2* (*33*) mice were obtained from S.Aizawa (RIKEN). *K14-H2BmCherry*(*34*) mice were generated in the laboratory and described previously. To visualize clonally labelled basal cells as they delaminate, *K14CreER; mTmG* mice were given a single dose of tamoxifen (20µg/g in corn oil) 3 days before imaging. *IvlCreERT2; R26-Tomato* mice were treated with tamoxifen once at 8 weeks (0.2 mg/g in corn oil i.p.; Sigma Aldrich, Cat#T5648), two days prior to cell isolation. To visualize K10 expression, *K10-rtTA*; *pTRE-H2BGFP*; *mTmG* mice were given doxycycline (2 mg/ml) in drinking water with 1% sucrose continuously, starting 3 days before imaging. To block proliferation in the K10 positive population, *K10-rtTA*; *pTRE-H2BGFP*; *mTmG; tetO-Cdkn1b* mice and *K10-rtTA*; *pTRE-H2BGFP*; *mTmG* littermate controls were given doxycycline (2 mg/ml) in drinking water with 1% sucrose continuously for the times specified. Mice from experimental and control groups were randomly selected from either sex for live imaging experiments. No blinding was done. All procedures involving animal subjects were performed under the approval of the Institutional Animal Care and Use Committee (IACUC) of the Yale School of Medicine or the Linköping Animal Ethics Committee in accordance to Swedish legislation.

#### *In vivo* imaging

Preparation of the skin and live imaging methods were similar to those performed previously(*10*, *23*). All imaging was performed in distal regions of the ear skin during prolonged telogen, with hair removed using depilatory cream (Nair) ≥3 days before the start of each experiment. Mice were anesthetized IP injection of ketamine/xylazine (15mg/ml and 1mg/ml, respectively in PBS) and anesthesia was maintained throughout the course of imaging with vaporized isofluorane delivered by a nose cone. Image stacks were acquired with a LaVision TriM Scope II (LaVision Biotec, Germany) laser scanning microscope equipped with both a Chameleon Vision II and Discovery 2-photon lasers (Coherent, USA). For collection of serial optical sections, the laser beam was focused through a 40x water immersion lens (Nikon; N.A. 1.15) and scanned with a field of view of 0.3mm x 0.3mm at 600Hz. Z-stacks were acquired with 0.5-1µm steps to image a total depth of ~40µm of tissue, covering the entire thickness of the epidermis. Visualization of ECM was achieved via second harmonic signal using blue channel at 940 nm imaging wavelength. To follow the same epidermal cells over multiple days, inherent landmarks of the skin together with a micro-tattoo were used to navigate back to the same epidermal regions every 12 or 24h. For time-lapse imaging, serial optical sections were obtained in a range of 5-8 minute intervals for a total duration of 1-3h.

#### Image Analysis

For quantifications in Figures 1, 3, and 4C-H, raw image stacks were imported into Fiji(*35*, *36*) for further analysis. Basal footprint and suprabasal spreading were quantified by manually outlining cell boundaries directly above the ECM signal and at the widest section in the top half of each cell, respectively. K10 reporter levels were quantified by measuring average H2BGFP signal intensity at the midpoint of each nucleus. K10 protein levels in whole mount images were quantified by measuring average cytoplasmic signal intensity at the midpoint of each cell. Cell behaviors were tracked by visually comparing epidermal regions at subsequent timepoints and cells were scored as suprabasal at the first timepoint when they made no observable contact with the underlying ECM signal. Prism software (Graphpad) was used to graph data and perform statistical analysis.

To track dynamics in the basal layer for Figure 4B, we adapted the procedure from(*23*). To height correct the 3D images, we first Gaussian blurred the signal from the ECM (mouse 1) or the epidermal cell nuclei (K14H2BmCherry) (mouse 2) spatially in the xy-plane (width 6 um) to create a 3D mask representing the region covering the whole epidermis. We then defined the height of the interface between the epidermis and the dermis from the 3D mask and subtracted this height from the original 3D data to level the basal layer position. From the height-corrected 3D images, we took three consecutive z-positions containing the nuclei of all the basal layer cells and averaged the intensity over the three slices to obtain 2D images in each channel. We calculated the local minima or maxima of the epithelial cell membranes (mem-tdTomato) or epithelial cell nuclei (K14H2BmCherry) to represent the cell positions in mouse 1 or mouse 2, respectively, and automatically corrected the shifts between time frames by minimizing the square distance between the nearest cell positions across the frames. The intensity of Keratin 10 reporter in the nucleus was calculated by taking the mean of the H2BGFP signal within a circle of 3 μm radius around the cell position. We manually set a threshold in the Keratin 10 signal level to define cell divisions with high Keratin 10 (22 out of 49 cell divisions in mouse 1, 46 out of 213 cell divisions in mouse 2).

To asses neighbor fate imbalance before cell division, we took the sum over the net imbalance of the fates in the frames leading up to each division event. The net imbalance was calculated by taking the time point right after the division as zero, and going back in time while adding one (differentiation) or subtracting one (division) whenever there was a fate event within in the 15 μm neighborhood of the division event of interest. We then averaged this net imbalance track over all the division events in mice 1 and 2. The error bar represents the fluctuation of net imbalance over each division event.

**Whole mount staining**

To isolate epidermis for staining, ear tissue was incubated in 5mg/ml dispase II solution (Sigma, 4942078001) at 37 °C for 10 minutes and the epidermis was separated from dermis using forceps. Epidermal tissue was fixed in 4 % paraformaldehyde in PBS for 10 minutes at room temperature, washed in PBS, permeabilized and blocked for >1h (0.2% Triton-X, 5% Normal Donkey Serum, 1% BSA in PBS) and then incubated in primary antibodies overnight at 4C and secondary antibodies for 3h at room temperature. Primary antibodies used were as follows: rabbit anti-K10 (1:1000; Biolegend Poly19054), guinea pig anti-K10 (1:400; Progen GP-K10), rabbit anti-pH3 (1:1000; Millipore 06-570), and chicken anti-GFP (1:1000; Invitrogen A10262). All secondary antibodies used were raised in a donkey host and were conjugated to AlexaFluor 488, 568 or 647 (Thermofisher). When used, AlexaFluor 647 Phalloidin (Thermofisher) was incubated at the same time as secondary antibodies. EdU was administered via intraperitoneal injection (50μg/g in PBS) 2h before harvesting tissue, and EdU labelling was performing using the Click-iT AlexaFluor 568 kit (Thermofisher) according to the manufacturer's instructions.

Fixed tissue was mounted on a slide with Vectashield Anti-fade mounting medium (Vector Laboratories) with a #1.5 coverslip.

**Fluorescent in situ hybridization (FISH)**

Single-molecule RNA-FISH (smRNA-FISH) was performed using the RNAscope Multiplex Fluorescent Detection Kit v2 (323100, Advanced Cell Diagnostics) according to manufacturer's instructions using TSA with Cy3, Cy5, and/or Fluorescein (NEL760001KT, Perkin Elmer) on FFPE sections of dorsal skin. FFPE sections were hybridized with combinations of the following mRNA probes (all from Advanced Cell Diagnostics): Krt10 (457901-C2), Krt14 (422521-C3), Krt5 (547901), Mt4 (447121-C3) and Krtdap (500671), together with cell membrane counterstaining using WGA (1:50, 29028-1, Biotium). RFP IHC (1:100, 600-401-379, Rockland) was performed together with smRNA-FISH stainings according to manufacturer's instructions. Each individual staining was performed on skin samples isolated from the same mice that were used for ITGA6-sorted cell sequencing as well as wt 8-week-old mice. Images were acquired on a Nikon A1R spinning disk confocal as tiled images (10%–15% overlap) and stitched by NIS Elements. Subsequently, all images were processed in the same way (maximum intensity projection, brightness adjustment, pseudocoloring) using Fiji(*35*, *36*).

**Single-cell RNA-sequencing and library preparation**

For the single-cell RNA-sequencing, epidermal cells were isolated from the back skin of 8-week-old *Ivl*-traced mice as described previously(*20*). Cells were stained for 1h with Cd49f (Itga6)-AlexaFluor 647 (1:50; BD Biosciences; Cat#551129), Sca1-PeCy7 (1:50; BD Biosciences; Cat#558162), and Cd34-FITC (1:50; BD Pharmingen Cat# 553733) and Sytox blue (1:1000; Life Technologies Cat# S34857) was added just prior (2 min before) to FACS sorting. Tomatotraced and non-traced live cells gated for ITGA6+/SCA1+/CD34-neg were collected in 400 µl Defined Keratinocyte serum-free medium (DK-SFM; Thermo Fisher, Cat#10744019) with 1% DNaseI (Stem Cell Technologies, Cat#07900), loaded into Fluidigm C1 chips (Fluidigm, USA) for library preparation and sequenced as described previously(*20*).

**Data analysis**

Pre-processing of the sequencing results into count matrices was performed as in(*20*), all subsequent data analysis was performed using Scanpy(*37*). From the previously published datasets (*20*, *38*) only cells that were categorized as "interfollicular epidermis cells" (IFE) were included. Each dataset (ITGA6-sorted, Joost 2016, Joost 2020) was separately normalized using size-factors and logaritmitized (ln(X+1)), before filtering out non-expressed genes. Further, each dataset was regressed for the effects of total counts per cell, percentage of ERCC spike-in counts for the ITGA6-sorted and Joost 2016 datasets (Fluidigm C1 based), and cell cycle (as scored by 'score_genes_cell_cycle' method). To combine all data, we first merged the C1-based datasets using the shared genes from the top 4000 highly variable genes from each individual dataset. For integrating the Joost 2020 dataset, we used genes present in the C1-merged data. To compensate for the lower per-cell read counts in the Joost 2020 dataset (10X Chromium based), Scanpy implementation of MAGIC imputation(*39*) was used with the following parameters specified: knn = 10; t = 2. This merged and imputed dataset revealed a small outlying group of cells (n=9 cells) and a population of infundibulum cells (n=147 cells), which were removed from subsequent analysis (see iPyhon notebook). Cell cluster classification was performed on the imputed data using the previously defined differentiation trajectory from Joost 2016 as a reference. A kNN-classifier (Scikit-learn(*40*)) was used to assign the cluster identity of cells from the ITGA6-sorted and Joost 2020 datasets according to the reference dataset with the following parameters specified: knn = 20; weight = 'distance'. For gene expression analyses, raw count matrices were log-normalized and downsampled to 2000 counts per cell, compensating for the higher read counts in Fluidigm C1-based datasets, and merged. Pseudotime analysis was performed using diffusion pseudotime implementation in Scanpy(*37*, *41*) and by ordering cells based on their position along the pseudotime. The cutoff for basal and suprabasal populations was defined by the 95$^{th}$ percentile of ITGA6-sorted cells on the ordered differentiation pseudotime. The cutoff for cells to be classified as *Krt10* positive was defined by the mean expression of *Krt10* in all cells (1.84 log-normalized counts). For further analysis of gene expression over pseudotime, cells were grouped into 6 and 4 equally sized bins for basal and suprabasal populations, respectively. Fitted gene expression trends along the pseudotime were compared to the average expression levels in bin 1 (considered to be the most basal) and plotted as the change (log-normalized) compared to this baseline expression. To avoid any bias due to sequencing-methods used and by merging the datasets, cell cycle analysis was performed on the largest dataset (before regression), covering males and females from different ages (Joost et al., 2020). Cell cycle phase was assigned with 'score_genes_cell_cycle' using a cutoff of 0.05 for positive classification. For mapping the (Aragona et al., 2020) dataset onto our differentiation timeline, only those cells defined in Aragona et al. as "Basal IFE", "Cycling IFE" and "Supra-basal IFE" were included (log-normalized and subset to include only genes that are expressed in all datasets). Finally, each dataset was independently scaled to unit variance and zero mean before mapping the Aragona et al. dataset to our combined UMAP using ingest implementation in Scanpy.

**Statistics and reproducibility**

Statistical parameters including the exact value of n for each experiment and statistical significance are reported in Figure Legends. Significance was determined using an unpaired Student's t-test or pairwise Tukey's HSD when multiple comparisons were made. Asterisks denote statistical significance (* $p<0.05$, ** $p<0.01$, *** $p<0.001$ and **** $p<0.0001$). Statistical calculations were performed using the Prism software package (GraphPad, USA).

**Supplementary Figure legends:**

**Supplementary Figure 1. Basal cell delamination lacks hallmarks of rapid, actomyosin-based extrusion. (A)** Cartoon schematic of epidermal structure. Stem cells reside in an underlying basal layer (directly above dermal extracellular matrix, shown in blue), and differentiate upwards to contribute to the outer barrier layers of the skin. **(B)** Current models of epidermal homeostasis propose that the basal layer is composed of either multiple distinct stem/progenitor cells (one common model proposes slow cycling stem cells that give rise to a self-sustaining population of differentiation-primed progenitors; left cartoon) or a single type of epidermal progenitor (right cartoon). **(C)** Representative images of a membrane GFP (*K14CreER*; *mTmG*) labelled cell revisted every 12h in the days preceding its exit from the basal layer. Top row: xy section from the widest point in the upper half of the cell. Middle row: xy section from directly above the ECM (blue). Bottom row: lateral reslice. Scale bar=10μm. **(D)** Quantification of basal cell-ECM contact in the days preceding basal layer exit. N=72 cells from 3 mice. ANOVA, $p<0.0001$; 36h vs 24h, 24h vs 12h, and 12h vs 0h Tukey's HSD, $p<0.0001$. **(E)** Quantification of suprabasal spreading, measured as the widest section in the upper half of each cell, in the days preceding basal layer exit. N=68 cells from 3 mice. ANOVA, $p<0.0001$; 36h vs 24h, 24h vs 12h, and 12h vs 0h Tukey's HSD, $p<0.0001$. **(F)** Representative image of Lifeact-GFP fluorescence (green) in the basal layer. Insets show a cell in the process of delaminating (arrows). Top inset is an xy section from the upper half of the cell; middle inset is an xy section from directly above the ECM signal (blue); bottom inset is a lateral reslice. **(G)** Representative image of GFP-tagged Non-muscle myosin IIB (*GFP-NMMIIB*) (green) in the basal layer. K14H2B-mCherry signal (red) shows positions of epithelial nuclei. Insets show a cell in the process of delaminating (arrows). Top inset is an xy section from the upper half of the cell, containing the cell nucleus; middle inset is an xy section from directly above the ECM signal (blue); bottom inset is a lateral reslice.

**Supplementary Figure 2. Merging scRNA-seq datasets.** **(A)** Representative sorting strategy with gates set for live (upper panel), ITGA6+ and SCA1+ (middle panel), and CD34- cells according to their *Ivl*-Tomato expression (lower panel). **(B)** Flowchart describing the workflow of merging ITGA6-sorted, Joost 2016 and Joost 2020 dorsal skin datasets. Grey lines indicate which steps were performed individually for each dataset and when they were combined. Gene counts for expression analysis were combined from log-normalized counts. **(C)** Locations of Joost 2016 cell clusters, overlaid on the combined UMAP, are colored according to local density of cells; all other cells are in grey. **(D)** Classification of cells in the combined dataset according to a kNN-classifier based on the reference clusters from Joost 2016: basal (B), differentiating (DI and DII), keratinizing (KI and KII) (left panel). Barplots showing the contribution of individual datasets to each cluster (right panel). (C-D) Dashed lines indicate the assigned basal-suprabasal border (delamination point).

**Supplementary Figure 3. Characterization of the scRNA-seq datasets.** **(A)** Classification of cells from all datasets as *Krt10*-positive or *Krt10*-negative (cutoff: 1.84 log-normalized and downsampled counts, see Methods). Cells are colored according to their *Krt10* expression levels. **(B)** Quantification of *Krt10*-positive and negative cells within the basal and suprabasal compartments (as defined by the delamination point). **(C)** Single-molecule RNA FISH validation of differentiation-associated gene expression in basal dorsal IFE cells. Upper panel: Evidence of *Krt10+*/*Krt14+* (yellow arrowheads) as well as *Krt10+*/*Krt14-* (white arrowheads) basal cells with a large ECM-footprint. Lower panel: Presence of a basal *Krt10+*/*Mt4+*/*Krtdap-* cell (arrowhead).

**(D)** Mapping of (Aragona et al., 2020) dorsal IFE dataset (colored) onto our combined UMAP (in grey). Colors indicate cluster annotations from Aragona dataset. **(E)** UMAP showing the grouping of cells into bins according to their location along the differentiation pseudotime. The basal-suprabasal border was set to be between bins 6 and 7 (Methods). **(F)** Changes in fitted gene expression levels (log-normalized) of the genes shown in Figure 2D, as compared to the baseline (average expression in bin 1). Solid and dashed colored lines indicate genes that respectively increase or decrease their expression as compared to bin 1. The basal-suprabasal border is between bin 6 and 7. **(G)** Expression patterns of genes shown in Figure 2D, overlaid on the combined UMAP. **(H)** Violinplots grouped according to pseudotime bins (left panel) and combined UMAPs (right panel) showing characteristic gene expression changes during the differentiation process. Expression levels are shown as log-normalized expression. (**I-J**) Single-molecule RNA FISH and antibody-based stainings showing basal *Ivl*-traced cells (Tom+ stained via RFP) with *Krt5* and *Mt4* co-expression (arrowheads) (I), and with *Krt14* and *Krt10* co-expression (arrowheads) (J). (C, I, J) WGA (wheat germ agglutinin) is used as a membrane stain. Dashed lines indicate the basement membrane. Scale bars: 25µm. (A, D, E, G, H) Dashed lines indicate the assigned basal-suprabasal border.

**Supplementary Figure 4. Characterization of Keratin 10-positive cell divisions. (A)** UMAP representation of different biological replicates of different ages from Joost 2020 dataset, overlaid on the merged dataset (in gray). **(B)** UMAP representation of cell cycle phases of Joost 2020 cells that were classified as *Krt10-positive*. **(C)** Quantification of the proportion of S and G2/M phase cells within the *Krt10-positive* cell population shown in (B). **(D)** Representative whole mount staining of pH3 (green) and K10 (red), showing both pH3-positive, K10 negative (yellow arrows) and pH3-positive, K10-positive (white arrows) basal cells. Scale bar=20μm. **(E)** Representative whole mount staining of mVenus-hGem signal from Fucci2 reporter mice (green), K10 (red) and phalloidin (white) showing both mVenus-hGem-positive, K10 negative (yellow arrows) and mVenus-hGem-positive, K10-positive (white arrows) basal cells. Scale bar=20μm. **(F)** Representative images of a revisited K10-positive basal cell as it divides to produce a delaminating daughter cell (number 1) and a daughter cell that goes through another round of cell division before exiting the basal layer (numbers 2a and 2b). Scale bar=10μm.

**Supplementary Figure 5. Details of a genetic approach to block proliferation in K10-positive basal cells. (A)** Schematic of strategy to block proliferation specifically in differentiating basal cells. K10 promoter-controlled rtTA was used to drive expression of both the G1 cell cycle inhibitor Cdkn1b (pTRE-Cdkn1b) and a fluorescent reporter (pTRE-H2BGFP) in a Doxycycline-inducible manner. **(B)** Quantification of K10 reporter positive mitotic figures after 24 hours of Doxycycline administration in Control and Cdkn1b mice. Student's t-test, $p<0.05$. **(C)** Quantification of delamination events (defined as the number of basal cells that lose ECM contact between Day 3 and Day 4 of Doxycyline administration) in Control and Cdkn1b mice. Student's t-test, $p>0.05$. For bar graphs in (B) and (C), error bars represent S.D.

**Supplementary Figure 6. A new model of epidermal basal layer heterogeneity.** We demonstrate that stem cells differentiate along a single, progressive continuum of transcriptional changes before they gradually delaminate out of the basal layer. Early transcriptional changes coincide with a commitment to differentiate but precede the loss of proliferative capacity, leading to distinct daughter cell fates depending on a cell's position along the trajectory.

**Supplementary Video 1. Mitotic division of K10 reporter positive basal cell.** Time-lapse recording over 88min of a representative K10 reporter positive basal cell dividing to produce two basal daughter cells. Top panel shows xz view and bottom panel shows xz view of the same cell. K10 reporter signal (*K10rtTA; pTRE-H2BGFP*) shown in green, cell membrane (mem-tdTomato) shown in red, dermal ECM shown in blue. Scale bar=5µm.

Figure S1

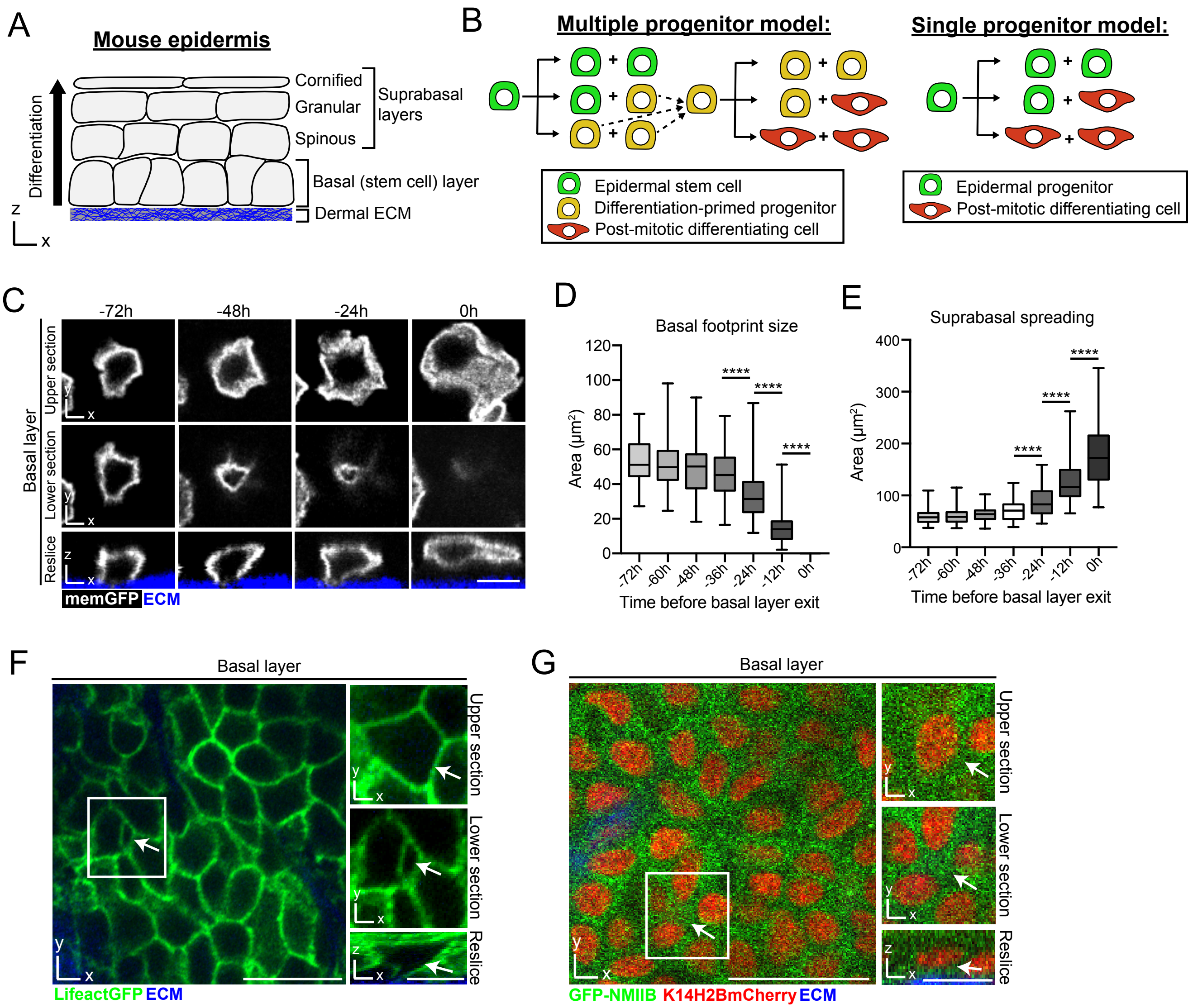
A
Mouse epidermis
Cornified
Granular
Spinous
Suprabasal layers
Basal (stem cell) layer
Dermal ECM
Differentiation
B
Multiple progenitor model:
Epidermal stem cell
Differentiation-primed progenitor
Post-mitotic differentiating cell
Single progenitor model:
Epidermal progenitor
Post-mitotic differentiating cell
C
-72h
-48h
-24h
0h
Basal layer
Upper section
Lower section
Reslice
memGFP
ECM
D
Basal footprint size
Area (µm²)
Time before basal layer exit
E
Suprabasal spreading
Area (µm²)
Time before basal layer exit
F
Basal layer
Upper section
Lower section
Reslice
LifeactGFP
ECM
G
Basal layer
Upper section
Lower section
Reslice
GFP-NMIIB
K14H2BmCherry
ECM

### Figure S2

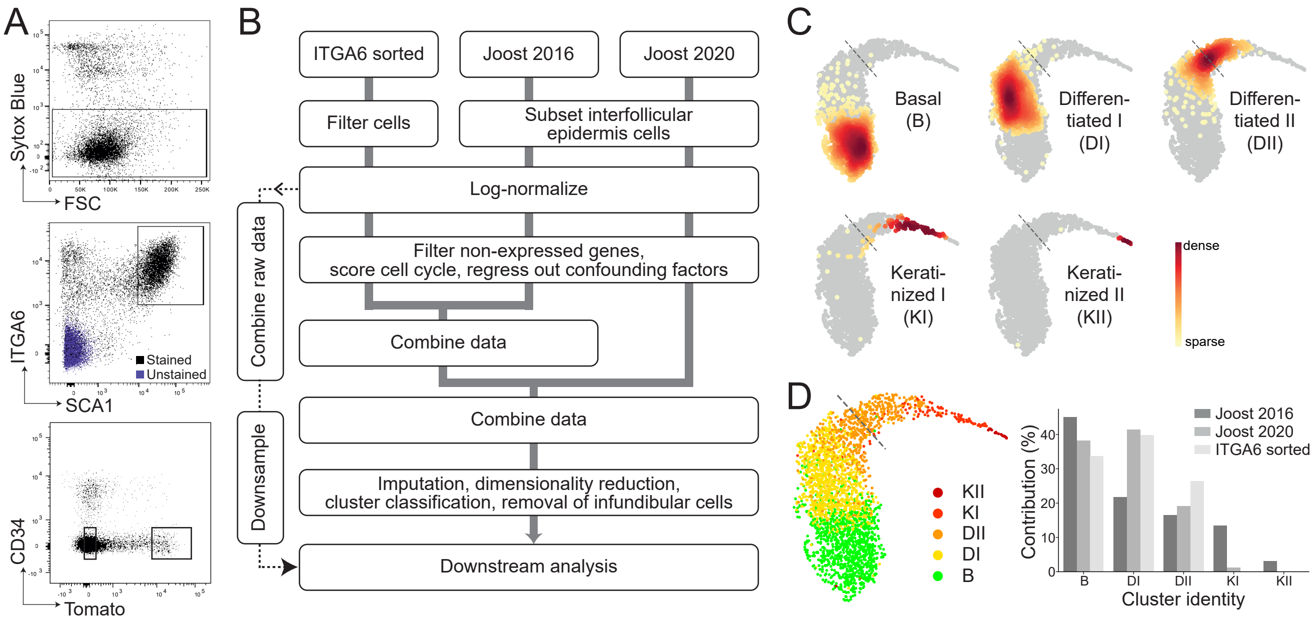

A
Sytox Blue
FSC
ITGA6
SCA1
Stained
Unstained
CD34
Tomato
B
ITGA6 sorted
Joost 2016
Joost 2020
Filter cells
Subset interfollicular epidermis cells
Log-normalize
Filter non-expressed genes, score cell cycle, regress out confounding factors
Combine data
Combine data
Imputation, dimensionality reduction, cluster classification, removal of infundibular cells
Downstream analysis
Combine raw data
Downsample
C
Basal (B)
Differen-tiated I (DI)
Differen-tiated II (DII)
Kerati-nized I (KI)
Kerati-nized II (KII)
dense
sparse
D
KII
KI
DII
DI
B
Joost 2016
Joost 2020
ITGA6 sorted
Contribution (%)
Cluster identity

Figure S3

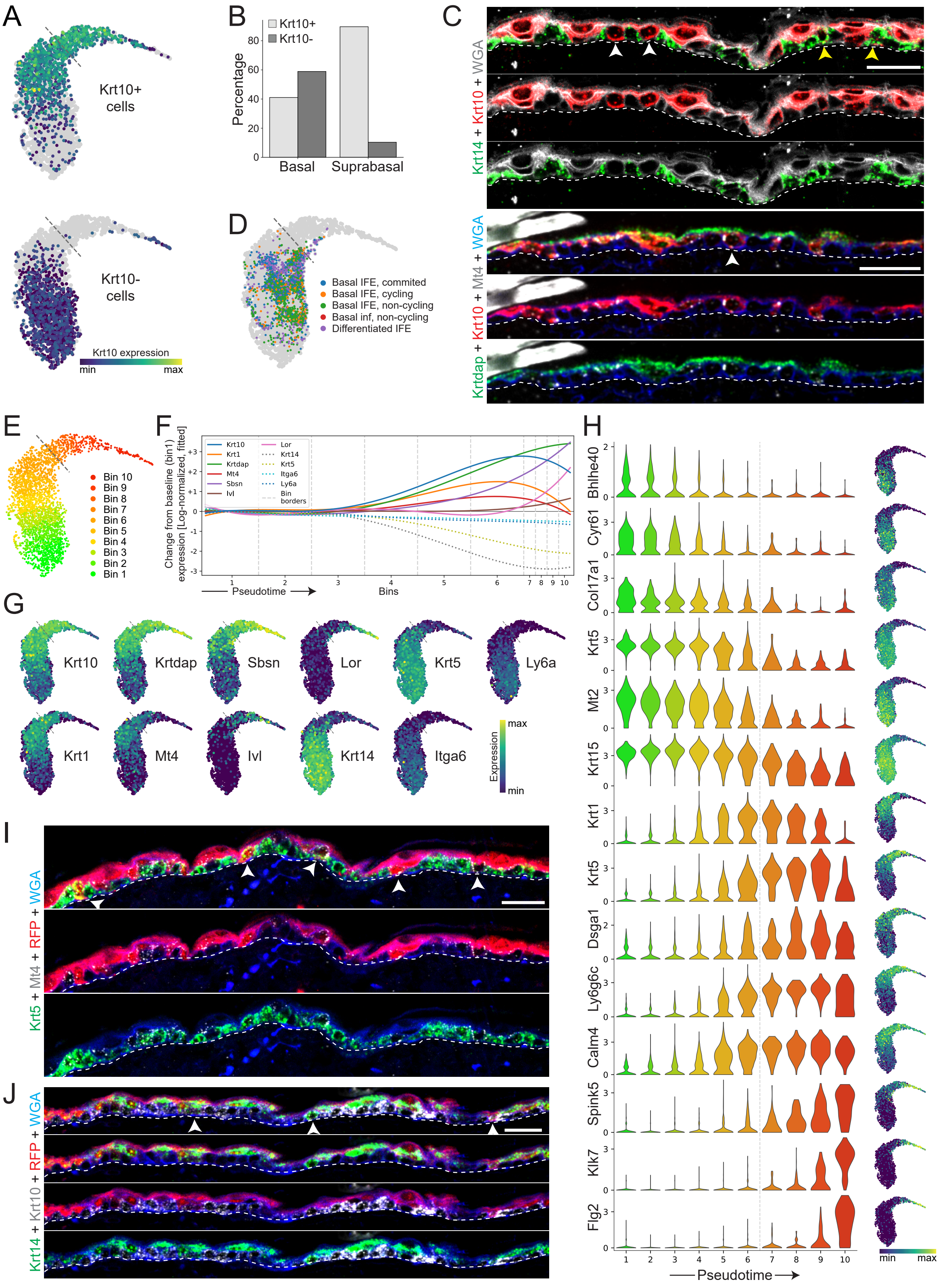

A
Krt10+ cells
Krt10- cells
Krt10 expression
min
max
B
Krt10+
Krt10-
Percentage
Basal
Suprabasal
C
Krt14 + Krt10 + WGA
Krtdap + Krt10 + Mt4 + WGA
D
Basal IFE, commited
Basal IFE, cycling
Basal IFE, non-cycling
Basal inf, non-cycling
Differentiated IFE
E
Bin 10
Bin 9
Bin 8
Bin 7
Bin 6
Bin 5
Bin 4
Bin 3
Bin 2
Bin 1
F
Change from baseline (bin1) expression [Log-normalized, fitted]
Krt10
Krt1
Krtdap
Mt4
Sbsn
Ivl
Lor
Krt14
Krt5
Itga6
Ly6a
Bin borders
Pseudotime
Bins
G
Krt10
Krtdap
Sbsn
Lor
Krt5
Ly6a
Krt1
Mt4
Ivl
Krt14
Itga6
Expression
max
min
H
Bhlhe40
Cyr61
Col17a1
Krt5
Mt2
Krt15
Krt1
Krt5
Dsga1
Ly6g6c
Calm4
Spink5
Klk7
Flg2
Pseudotime
min
max
I
Krt5 + Mt4 + RFP + WGA
J
Krt14 + Krt10 + RFP + WGA

### Figure S4

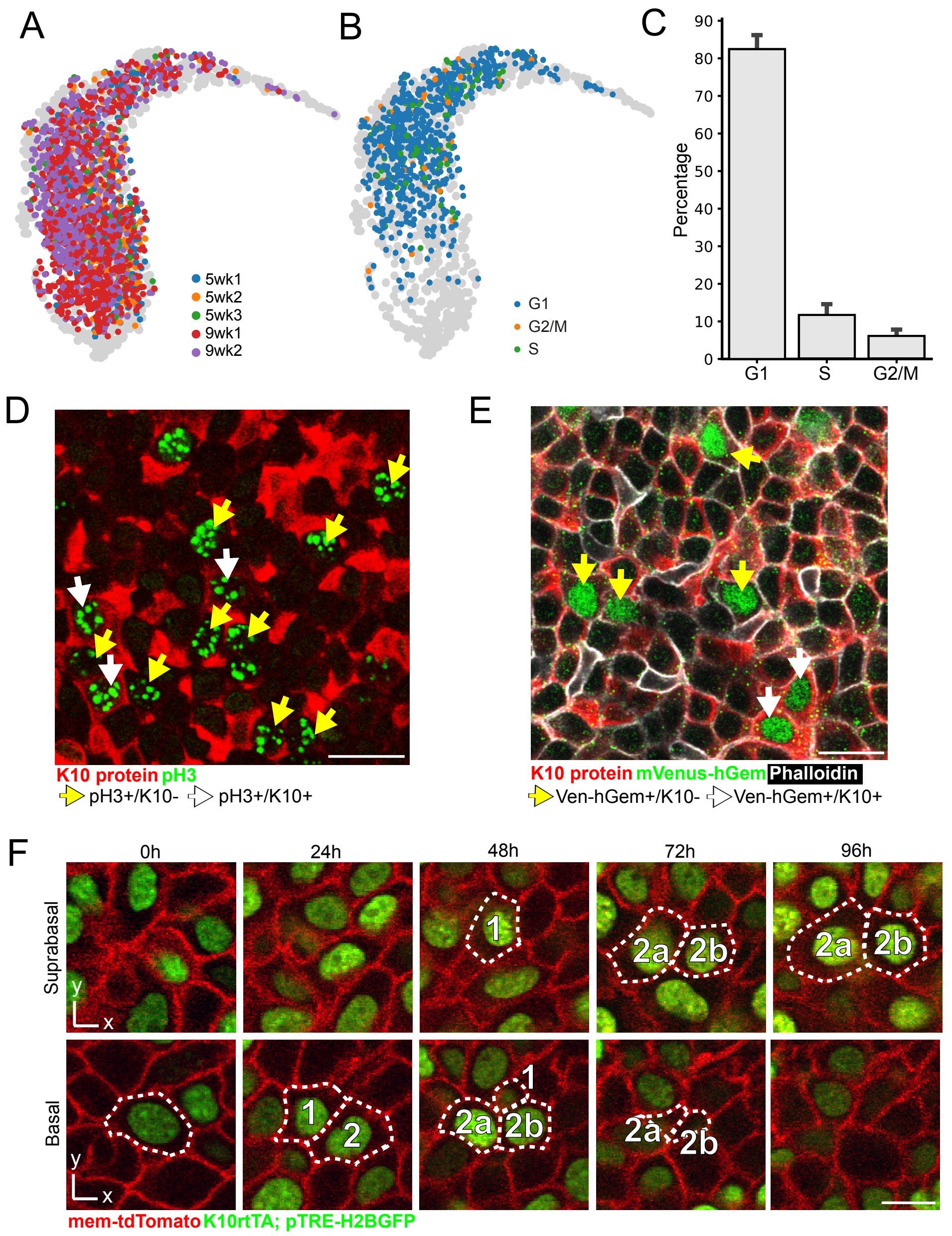
A
5wk1
5wk2
5wk3
9wk1
9wk2
B
G1
G2/M
S
C
Percentage
90
80
70
60
50
40
30
20
10
0
G1
S
G2/M
D
K10 protein pH3
pH3+/K10-
pH3+/K10+
E
K10 protein mVenus-hGem Phalloidin
Ven-hGem+/K10-
Ven-hGem+/K10+
F
0h
24h
48h
72h
96h
Suprabasal
Basal
y
x
1
2
2a
2b
mem-tdTomato K10rtTA; pTRE-H2BGFP

### Figure S5

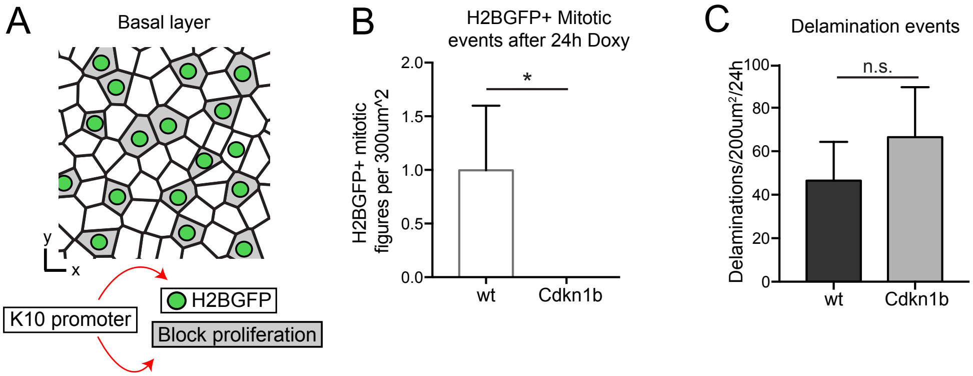

A
Basal layer
y
x
K10 promoter
H2BGFP
Block proliferation
B
H2BGFP+ Mitotic events after 24h Doxy
H2BGFP+ mitotic figures per 300um^2
2.0
1.5
1.0
0.5
0.0
*
wt
Cdkn1b
C
Delamination events
Delaminations/200um²/24h
100
80
60
40
20
0
n.s.
wt
Cdkn1b

### Figure S6

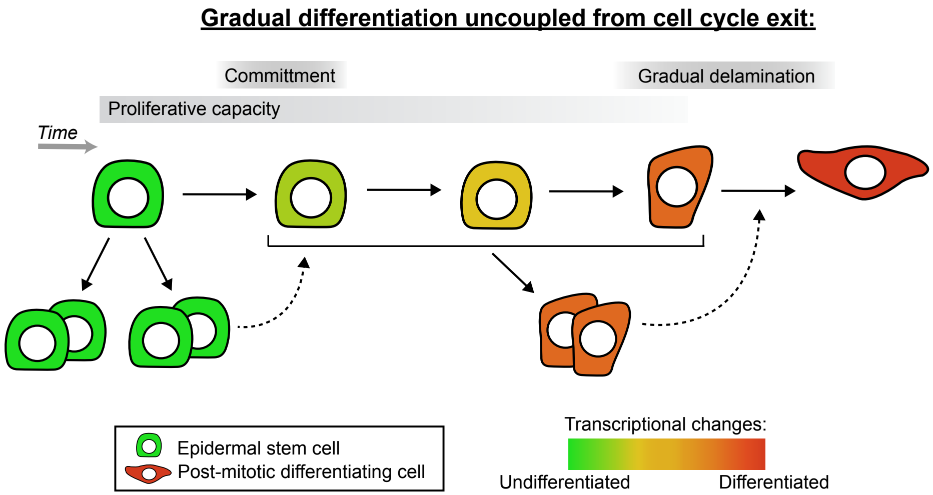
Gradual differentiation uncoupled from cell cycle exit:
Committment
Gradual delamination
Proliferative capacity
Time
Epidermal stem cell
Post-mitotic differentiating cell
Transcriptional changes:
Undifferentiated
Differentiated
